# Lectin-Seq: a method to profile lectin-microbe interactions in native communities

**DOI:** 10.1101/2022.12.14.520458

**Authors:** Robert L. McPherson, Christine R. Isabella, Rebecca L. Walker, Dallis Sergio, Tony Gaca, Smrithi Raman, Le Thanh Tu Nguyen, Darryl A. Wesener, Melanie Halim, Michael Wuo, Amanda Dugan, Robert Kerby, Soumi Ghosh, Federico E. Rey, Hera Vlamakis, Eric J. Alm, Ramnik J. Xavier, Laura L. Kiessling

## Abstract

Soluble human lectins are critical components of innate immunity. Genetic models suggest lectins influence host-resident microbiota, but their specificity for commensal and mutualist species is understudied. Elucidating lectins’ roles in regulating microbiota requires understanding which microbial species they bind within native communities. To profile human lectin recognition, we developed Lectin-Seq. We apply Lectin-Seq to human fecal microbiota using mannose-binding lectin (MBL) and intelectin-1 (hItln1). The microbial interactomes of MBL and hItln1 differ in composition and diversity. MBL binding is highly selective for a small subset of species commonly associated with humans. In contrast, hItln1’s interaction profile encompasses a broad range of lower-abundance species. Thus, human lectins have evolved to recognize distinct species of commensals, suggesting they directly influence microbiome composition. Lectin-Seq offers a new means of annotating microbial communities.

**One-Sentence Summary:** Soluble human lectins bind distinct bacterial species in fecal microbiota.

## Main text

Humans host trillions of microbes (*1*), but the mechanisms we use to recognize and distinguish between different microbial species are not fully defined. Microbes are decorated with cell surface glycans that act as a primary point of contact between host and microbe (*2*). Microbial glycan composition is unique between species and could serve as a molecular readout of cellular identity for host factors (*3*). Soluble human carbohydrate-binding proteins termed lectins distinguish cell surface glycans and can achieve species- and strain-level binding specificity (*4–6*). Lectins are implicated in immune defense against pathogens, and lectin-microbe interactions can be key determinants of cell fate for microbial pathogens (*7, 8*). Genetic models suggest lectins play significant roles in regulating broad groups of microbial species in the gut microbiota (*9–12*). However, the microbial specificity of human lectins in host-resident communities has not been elucidated. Critically, it is unclear whether lectins in such communities bind a wide range of species or restrict their interactions to select microbes. Such information is vital to developing a mechanistic understanding of how lectins regulate the microbiota.

Lectins are commonly classified by their monosaccharide specificity, but their cell binding specificity can be influenced by glycan accessibility, density and secondary interactions (*13*). In some cases, monosaccharide specificity can be used to predict lectin-microbe interactions (*14*), but we postulate that other features of cell surface glycan display influence recognition. Previous studies have primarily characterized lectin-microbe interactions using monoculture binding assays (*11, 14*). Single-species binding assays have drawbacks: bacteria can modulate their glycan displays under different growth conditions potentially leading to significant differences in the glycan composition of native and cultured microbes (*15, 16*). Moreover, relevant lectin binding occurs in mixed communities. We postulate that competition for lectin binding might occur in such physiologically relevant settings.

To assess lectin selectivity within mixed communities, we first evaluated the binding of the human lectin intelectin-1 (hItln1) to bacterial isolates from the human gut microbiota. Previously, we found hItln1 exhibits serotype-specific binding to isolates of *Streptococcus pneumoniae* (*14*). Others have noted binding to different bacterial species considered pathogenic, suggesting hItln1 targets pathogens (*17, 18*). Since hItln1 is expressed in the small intestine (*19*), we assessed the binding of recombinant hItln1 to a collection of human gut microbial strains by flow cytometry, and observed binding to strains from diverse phyla (Fig. 1A-B; Table S1). The ability of hItln1 to bind to gut-resident species that can contribute to human health emphasizes the need to further characterize human lectin binding to commensal and mutualist microbes.

**Figure 1.**
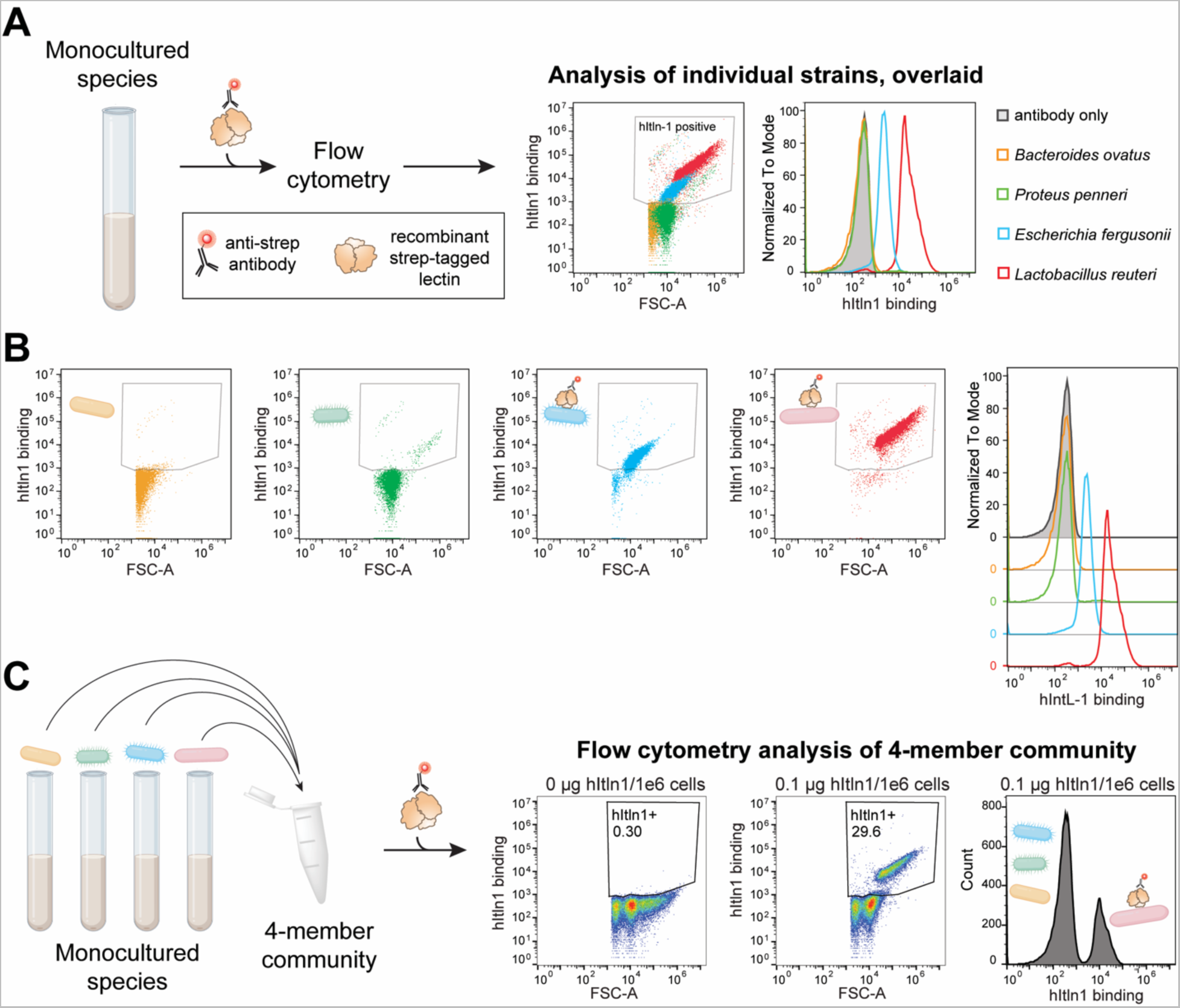
hItln1 binding in a synthetic mixed microbial community. (A) Schematic of flow cytometry-based assay to measure soluble lectin binding to microbe isolates. Overlay of data from 0.1 μg hItln1/1e6 cells condition shown in (B) with the hItln1-positive binding region indicated. (B) Flow cytometry analysis of microbe isolates plotted as forward scatter (FSC) against hItln1 binding (anti-Strep Oyster 645). Panels show 0.1 μg hItln1/1e6 cells condition. Staggered histograms of flow cytometry data plot cell count as a percent of the maximum against lectin binding (anti-Strep Oyster 645). Dot plots and histograms are color-coded according to the legend shown in (A). (C) Schematic of flow cytometry-based assay to measure soluble lectin binding to a synthetic mixture of microbe isolates. Flow cytometry analysis of synthetic mixed microbial community plotted as forward scatter (FSC) against hItln1 binding (anti-Strep Oyster 645). Panels show 0 μg hItln1/1e6 cells and 0.1 μg hItln1/1e6 cells conditions. Histogram of flow cytometry data plots cell count as a percent of the maximum against lectin binding (anti-Strep Oyster 645) for the 0.1 μg hItln1/1e6 cells condition.

We then generated a synthetic community from two binding species (*Escherichia fergusonii* and *Lactobacillus reuteri*) and two non-binding species (*Bacteroides ovatus* and *Proteus penneri*) that were each distinguishable by their scattering properties (Fig. 1A-B). In this mixed community of 50% hItln1 binders (Fig. 1C), the lectin bound to only 29.6% of cells. Even at higher hItln1 concentrations, 50% binding was never achieved (Fig. S1A). The forward scatter (FSC) profile of the hItln1-bound population matches that of *L. reuteri* (Fig. 1A-C, Fig. S1B), indicating that *L. reuteri* outcompetes *E. fergusonii* for htln1 binding in this simple mixed community. These data show that lectin selectivity can be altered in microbial mixtures, providing impetus to profile lectin binding to bacterial species within native communities.

We next characterized the binding specificity of two lectins in native microbial communities. The human gut harbors the most complex and diverse microbial community associated with the human body, and snapshots of the community are readily accessible through stool (*20*). We queried binding by hItln1, as its expression in the gut (*19*), its biochemical affinity for glycans exclusively found on microbes (*14, 21*), and our initial binding data (Fig. 1) all suggest it may preferentially interact with gut-resident microbes. Additionally, hItln1 is associated with diseases linked to microbiome dysbiosis, suggesting it may regulate gut microbiota (*22, 23*). However, interactions between hItln1 and intestinal microbial communities have not been assessed.

To contrast with our studies of hItln1, we also evaluated the specificity of human mannose-binding lectin (MBL) for human gut microbial communities. Decades of research show MBL binding promotes pathogen clearance (*24–26*) supporting a role in immune defense. A deficiency of MBL in murine models causes significant changes in the gut microbiota (*12*), but whether these changes are due to direct MBL-microbe interactions is unclear. MBL is primarily expressed in the liver, but low levels of expression have been detected in the intestine (*27*). An understanding of the direct interactions between MBL and gut-resident microbes is needed to interpret its influence on microbiome regulation.

We used flow cytometry to analyze the binding of recombinant MBL and hItln1 to stool homogenates collected from healthy human donors (Fig. 2A). In the presence of calcium ions, MBL and hItln1 each bound 15-20% of intact microbial cells in stool homogenate (Fig. 2B, middle column; Fig. 2C). Both lectins require calcium to bind glycans (*28, 29*), and analysis after the addition of the calcium chelator EDTA indicates the detected microbial binding interactions are glycan-dependent (Fig. 2B, right column). MBL and hItln1 bind orthogonal sets of glycans on a microbial glycan array (Fig. 2D; Table S2). The two lectins show similar levels of bacterial engagement (Fig. 2C), and we hypothesized their orthogonal glycan specificities would translate to specificity for different microbes. To this end, we co-incubated fluorescently labeled MBL and hItln1 with healthy human stool homogenate and analyzed binding by flow cytometry. Most lectin-bound cells were bound by either MBL or hItln1, while <5% of were co-bound by both lectins (Fig. 2E-F). Thus, MBL and hItln1 bind distinct bacterial populations.

**Figure 2.**
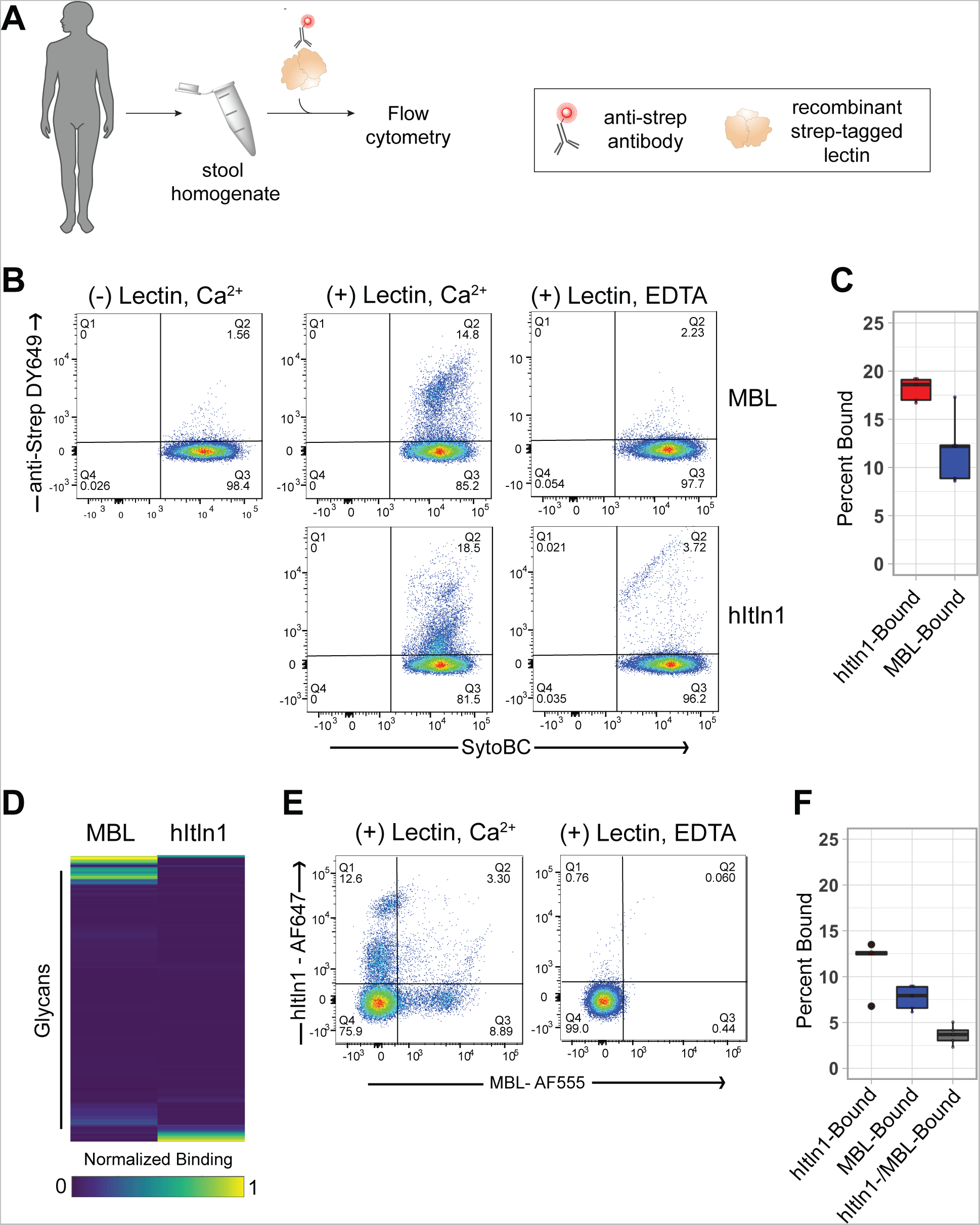
MBL and hItln1 binding to microbial populations in human stool samples. (**A**) Schematic of flow cytometry-based assay to measure soluble lectin binding to stool microbiome. (B) Flow cytometry analysis of healthy human stool plotted as SytoBC against lectin binding (anti-Strep DY649). Panels show no lectin control (column 1), and strep-MBL (top row) or strep-hItln1 (bottom row) binding in the presence of Ca^2+^ (column 2) or EDTA (column 3). (C) Quantification of data shown in A. Percent bound is calculated by determining the percentage of cells that fall in the strep-pos/SytoBC-pos fraction (Q2). Data shown are from the (+) strep-hItln1/Ca^2+^ (red; n=5; 18.1 +/- 0.5 % (mean, SEM)) and (+) strep-MBL/Ca^2+^ (blue; n = 5; 11.9 +/- 1.6%) samples. (D) Heatmap depicting normalized binding of MBL and hItln1 to microbial glycan array. Each row represents binding to a single glycan (mean of three spots). Rows were grouped by hierarchical clustering (Euclidean). (E) Flow cytometry analysis of the stool microbiome to determine differential hItln1 and MBL binding. Flow cytometry data is plotted as hItln1-binding (AF-647) against MBL-binding (AF-555). Panels show lectin-binding in the presence of (+) lectin/Ca^2+^ (panel 1) or (+) lectin/EDTA (panel 2). (F) Quantification of data shown in D, panel 2. Percent bound is calculated by determining the percentage of cells that fall in the hItln1-pos/MBL-neg (red; n = 5; 11.6 +/- 1.2%), hItln1-neg/MBL-pos (blue; n = 5; 7.7 +/- 0.6%) and hItln1-pos/MBL-pos fractions (dark grey; n = 5; 3.6 +/- 0.5%).

Our next objective was to identify the microbial taxa recognized by each lectin. To maximize microbial diversity in our analyses we included stool homogenate from 5 healthy and 9 clinically diagnosed IBD patient donors. While the composition of healthy and IBD microbiomes differ (*30*), we observed similar levels of lectin binding between healthy and diseased samples (Fig. S2A). Stool homogenates from these 14 donors were stained with MBL or hItln1, and the lectin-bound and unbound cells were isolated by fluorescence-activated cell sorting (FACS). The sorted fractions were analyzed by metagenomic sequencing and assigned species-level taxonomy using MetaPhlAn2.0 (*31*) to determine the relative abundance of microbial species in each fraction. Finally, we calculated a probability ratio (P_R_) (*32*) for each species to determine if it was enriched or depleted in the lectin-bound fraction. We refer to this workflow as Lectin-Seq (Fig. 3).

**Figure 3.**
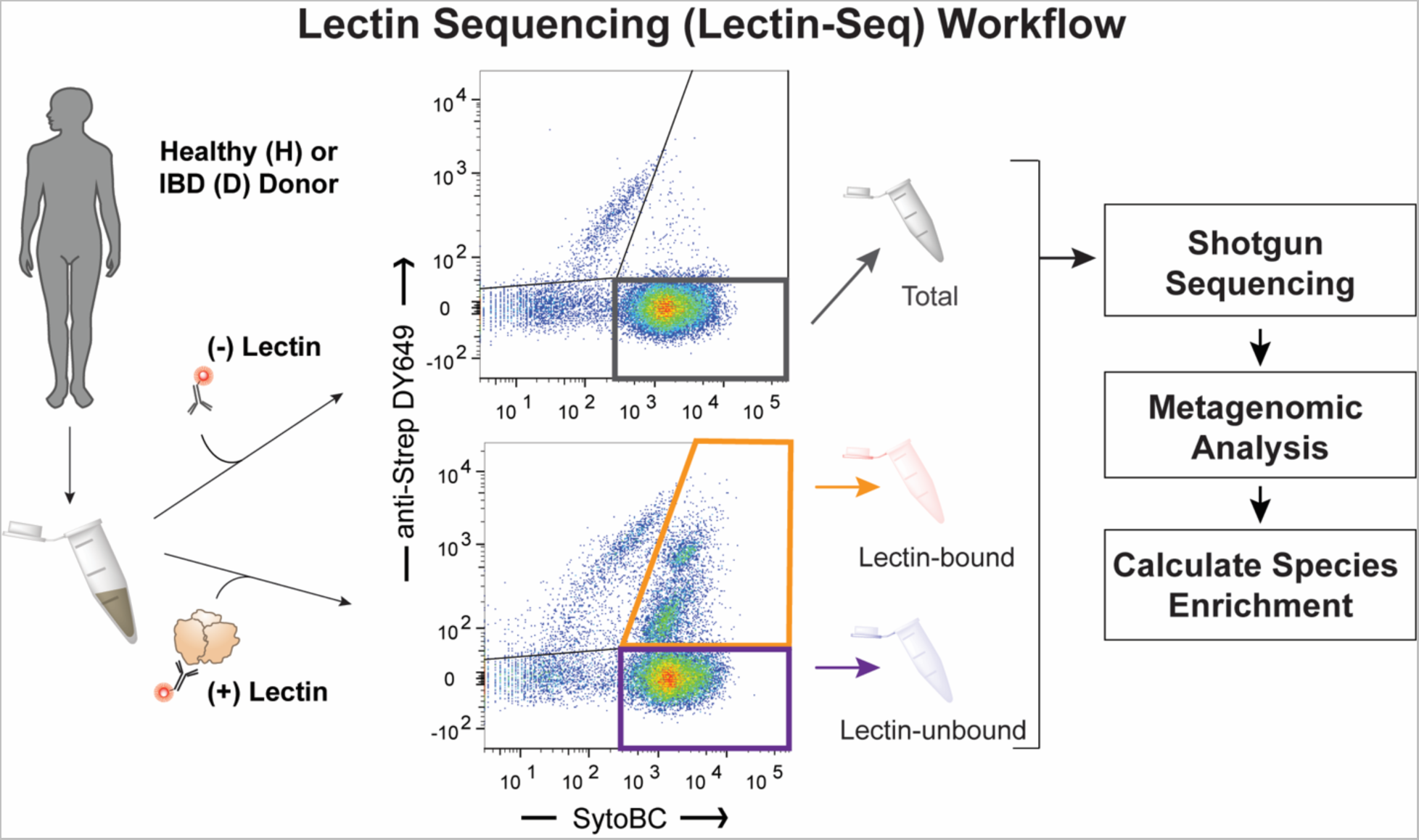
Lectin-Seq workflow for identifying lectin bound microbes from human stool samples.

The microbe-binding selectivity of MBL was remarkably specific. Lectin-Seq revealed that MBL-bound fractions were predominantly composed of *Faecalibacterium prausnitzii* (Fig. 4A, Fig. S3; Table S3). *F. prausnitzii* was present in the MBL-bound fractions of 12/14 stool samples and on average accounted for >40% relative abundance across all MBL-bound fractions (Fig. 4A-B; Fig. S3). In the two remaining samples, we did not detect *F. prausnitzii* in the input fraction (Fig. 4B). In stool samples that contain *F. prausnitzii*, the microbe was enriched in the MBL-bound fraction of 8/12 samples as indicated by a positive P_R_ (Fig. 4C). In the remaining 4 samples *F. prausnitzii* had a negative P_R_ close to zero indicating that this microbe was present at similar levels in the MBL-bound and unbound fraction (Fig. 4C).

**Figure 4.**
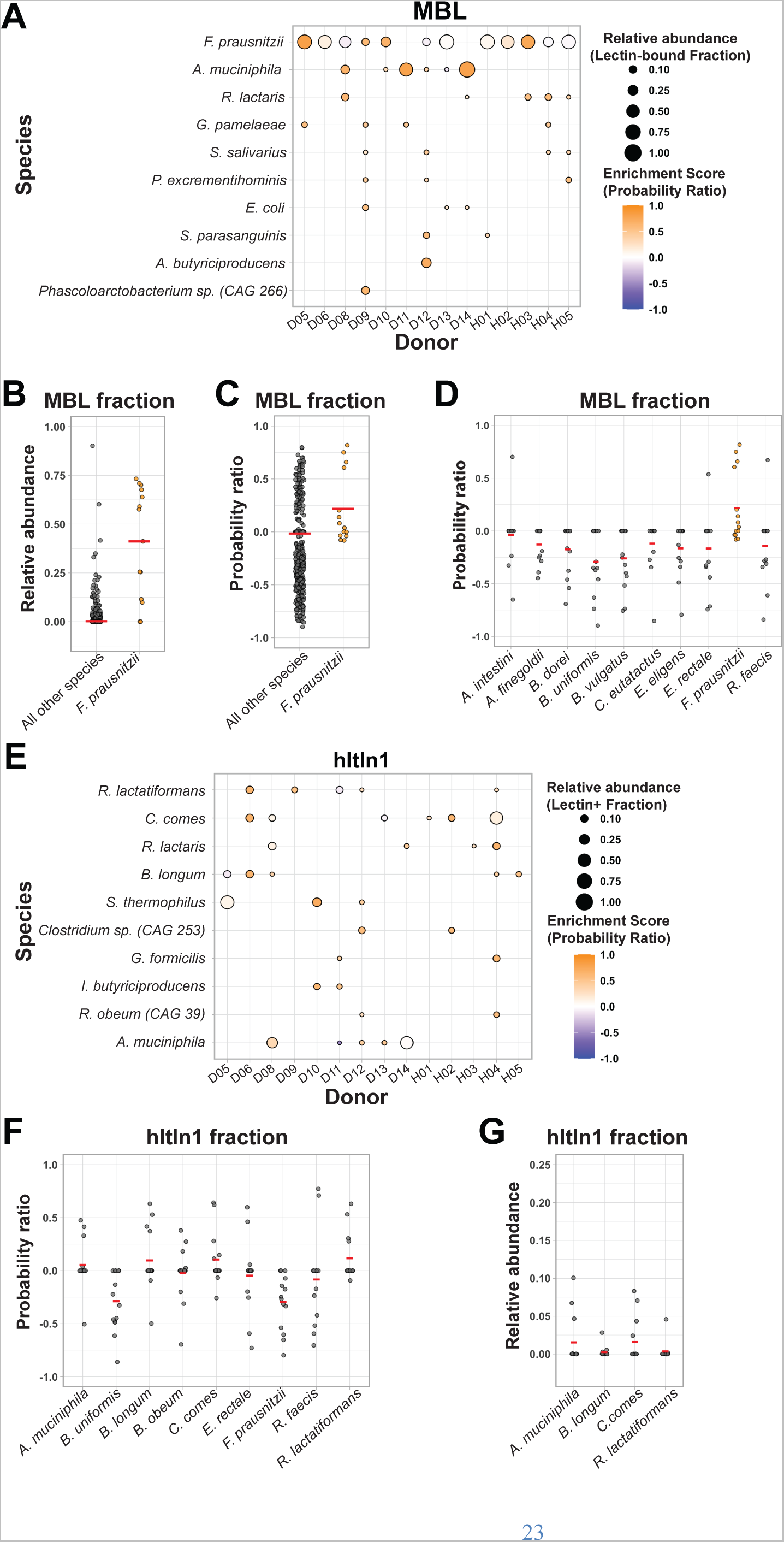
Identification and quantification of MBL and hItln1 bound bacterial species from human stool samples. (**A, E**) “Bubble plot” depicting the enrichment of top enriched bacterial species in the lectin-bound fraction (defined as the Probability Ratio) donor stool samples bound by either MBL (A) or hItln1 (E). Panels show top enriched bacterial species as calculated by average probability ratio. Probability ratio of each species is represented by the color of each dot, and fractional abundance is represented by the size of the dot. (B-C) Dot plot of the (B) relative abundance or (C) probability ratio of *F. prausnitzii* or all other species in the Lectin+ Fraction (MBL; all samples). Red line denotes the mean. (D) Dot plot of the probability ratio of top 10 most abundant species (input fraction) in the MBL+ fraction. Red line denotes the mean. (F) Dot plot of the probability ratio of species found in >5 hItln1+ fractions across samples. Red line denotes the mean. (G) Dot plot of the relative abundance (input fraction) of species with P_R_ > 0 (hItln1+ fraction). Red line denotes the mean.

*F. prausnitzii* is a highly abundant member of the gut microbiota; in the samples analyzed, the average percent relative abundance was ∼12% (Table S3). To determine if MBL recognition correlates with species abundance, we asked if the MBL-bound fraction was enriched for other abundant species. Out of the top 10 most abundant species (average abundance in input fraction; Fig. S4A) only *F. prausnitzii* had an average P_R_ > 0 for the MBL-bound fraction, indicating that MBL binding does not track with species abundance (Fig. 4D). MBL enriched for an additional 35 lower relative abundance species (average input abundance <0.01); however, >50% of these species are found in only one stool sample (Table S3; Fig. S5A).

**Figure 5.**
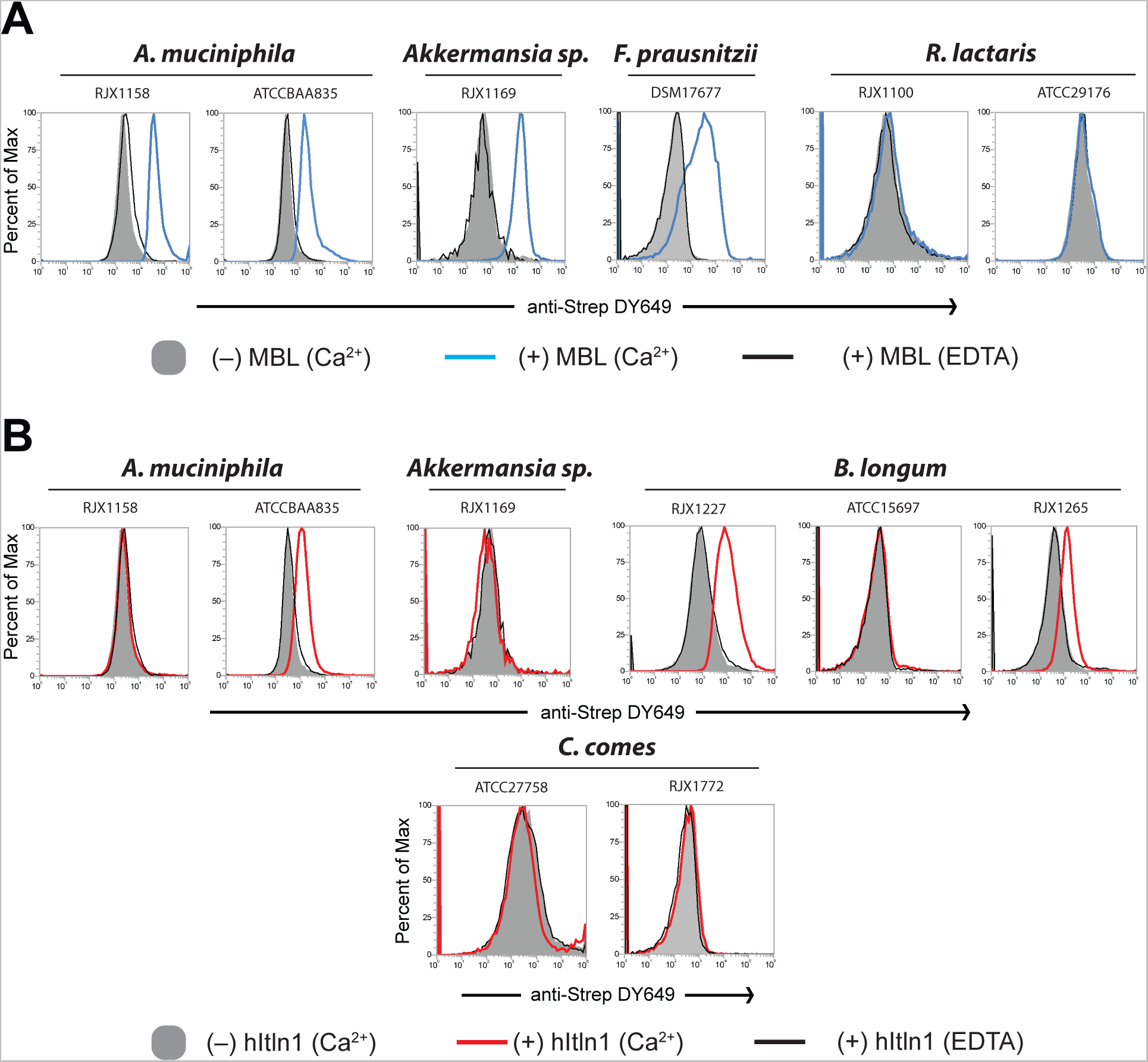
MBL- and hItln1-binding to bacterial isolates. (A-B) Summarized flow cytometry analyses of MBL (A) or hItln1 (B) binding to bacterial isolates. Isolates are grouped by species. Individual histograms plot cell counts as a percent of the maximum signal against lectin binding (anti-Strep DY649). Each plot shows data from (-) lectin (Ca^2+^; solid grey), (+) MBL/Ca^2+^ (blue trace) or hItln1/Ca^2+^ (red trace), and (+) lectin/EDTA (black trace) binding conditions. Data are representative of two independent experiments.

There were only 2 species apart from *F. prausnitzii* that were enriched in the MBL-bound fractions of at least five samples – *Akkermansia muciniphila* and *Ruminococcus lactaris* (Fig. 4A; Table S3; Fig. S5A). Notably, MBL binding to *A. muciniphila* was detected only in IBD-diagnosed donor samples including the 2 samples that lacked *F. prausnitzii* (Fig. 4A; Fig. S4B-C). *F. prausnitzii* levels are reduced in IBD patients (*33*); therefore, MBL should exhibit altered or broadened selectivity when its preferred binding partner is absent or present at lower levels. Additionally, *A. muciniphila* isolates from different individuals are genetically and phenotypically diverse (*34, 35*). Thus, *A. muciniphila* strains associated with disease may have an altered glycan display that leads to differential MBL binding.

MBL and hItln1 bound a similar absolute proportion of cells in samples, but the taxa bound by hItln1 and MBL did not significantly overlap. While MBL preferred *F. prausnitzii*, hItln1-bound fractions consisted of more diverse populations (Fig. S3; Fig. S5B; Table S3). No single species accounted for >15% of the average % relative abundance in hItln1-bound fractions (Fig. 4E; Fig. S3; Fig. S5B; Table S3), and the targeted species were not found in most of the enriched fractions. Nine species were found in five or more hItln1-bound fractions; however, the P_R_ values for these species varied across samples indicating that enrichment by hItln1 is donor dependent (Fig. 4F; Fig. S5B). Four species – *A. muciniphila*, *Bifidobacterium longum*, *Ruthenibacterium lactatiformans* and *Coprococcus comes* – had an average positive P_R_ across all samples (Fig. 4F). These four species each made up less than 2% of the input across all samples and were not detected in the input fractions of most donor samples (Fig. 4G). These findings indicate that hItln1 binds to low-abundance species within the human microbiome, while MBL prefers *F. prausnitzii*, a common and prevalent commensal.

To compare Lectin-Seq to *in vitro* strain binding, we tested MBL and hItln1 binding to select isolates of highly enriched species. MBL bound all tested strains of *A. muciniphila* and *F. prausnitzii* supporting the robust enrichment of these species we observed using Lectin-Seq (Fig. 5A). We also verified that MBL and hItln1 did not bind isolates of species that were depleted from the respective lectin-bound fractions in stool homogenate (Fig. S6). Intriguingly, however, we observed that some interactions are strain specific. For example, MBL did not bind any of the tested isolates of *R. lactaris* (Fig. 5A) despite its ability to enrich for this species from human samples. This attribute of the lectins was even more pronounced for hItln1, which bound 2/3 isolates of *B. longum*, 1/2 isolates of *A. muciniphila*, and no isolates of *C. comes* (Fig. 5B).

Because strain identity varies across individual microbiotas (*36*), the donor-dependent enrichment of these species by hItln1 may be attributed to strain-specific lectin interactions. These findings highlight the importance of evaluating lectin binding to endogenous bacterial populations. Our data support our findings that MBL binds *F. prausnitzii* and *A. muciniphila* in human samples and provide a mechanistic explanation for the donor-dependent interaction profile of hItln1.

Historically, the physiological roles of soluble lectins are ascribed to pathogen recognition and clearance (*6*), but emerging data suggest a role for murine lectins in regulating gut microbiota (*9–12*). Our findings provide direct evidence that human lectins bind selectively to distinct bacterial species in native communities. Lectin-Seq showed MBL robustly binds *F. prausnitzii*, an abundant member of the gut microbiome, and to a lesser degree *A. muciniphila* and *R. lactaris*. Such specificity suggests MBL could be co-opted as a tool to rapidly quantify the abundance of these species in uncharacterized stool samples. The specificity of MBL binding to mutualist and commensal species contrasts with previous data indicating that MBL exhibits relatively indiscriminate binding to a broad range of pathogenic microbes (*37*).

The high specificity of MBL binding to host-resident microbes suggests its biological functions go beyond pathogen surveillance. MBL has been reported to be expressed at low levels in the intestine (*27*). MBL could specifically bind to a proportion of abundant commensals such as *F. prausnitzii* to limit their growth and allow for propagation of lower-abundance species. *Akkermansia* levels are decreased in MBL knockout mice (*12*), but it is unclear if this is due to direct interactions of these microbes with murine MBL or if this finding translates to humans. Further mechanistic work is needed to determine the effect of MBL binding on the growth and colonization of these various microbial binding partners in the human gut. In the blood, MBL binding to pathogens results in their clearance (*24*); therefore, in healthy individuals MBL may be poised to eliminate specific gut-resident microbes that breach epithelial barriers and reach the blood. Conversely, in patients diagnosed with IBD or colorectal cancer (CRC) a reduction in barrier integrity can lead to blood, and potentially high levels of MBL, entering different regions of the gut. Indeed, *F. prausnitzii* levels are broadly reduced in patients with IBD (*33*) and, more specifically, in CRC patients that report blood in their stool (*38*). The binding of MBL to *F. prausnitzii* is consistent with these observations.

Lectin-Seq revealed that hItln1 exhibits donor-dependent binding to a broad group of low abundance microbes. hItln1-bound microbes are highly diverse: the population includes Gram-positive and -negative species capable of critical metabolic activities such as short-chain fatty acid production and mucin degradation. The implications of hItln1 binding to these gut-resident species are unclear. The interaction of hItln1 with select pathogens activates macrophage- and neutrophil-mediated killing (*17, 18, 39*); however, hItln1-mediated neutrophil activation requires serum components suggesting the bactericidal functions of hItln1 may be limited to the blood. Unlike some intestinal lectins (*11*), hItln1 has no known microbial toxicity and, unlike MBL, lacks canonical domains for complement recruitment (*14*). Alternatively, hItln1 is reported to suppress lipopolysaccharide-induced macrophage activation (*40, 41*), suggesting that under specific conditions hItln1 could play a protective role to shield microbes from macrophage killing. Thus, the function of hItln1 in innate immunity appears to depend on context. Further examination of hItln1 functions in the GI immune response is needed to determine whether hItln1 acts in a bactericidal capacity to clear specific gut-resident microbes from the body, or in a protective role to shield diverse groups of low abundance species from immune clearance and promote their colonization of the gut.

Our data provide insight into the interplay of lectin recognition and microbial glycans. A major finding from our study is that soluble human lectins have strikingly disparate microbial interaction profiles. MBL is highly specific for a prevalent human gut commensal, while hItln1 binds a diverse group of low-abundance species. Further, the orthogonal glycan specificities of these lectins reflect the importance of microbial glycans for species-specific host-microbe interactions. Still unclear is whether MBL and hItln1 have evolved to recognize ligands differentially displayed across the microbiota, or if specific members of the microbiota have evolved mechanisms allowing them to adjust their glycan displays to engage or evade these lectins. Future efforts to identify the microbial genetic determinants of lectin binding will inform the mechanistic basis and evolutionary dynamics of these host-microbe interactions. Our data highlight the utility of Lectin-Seq for uncovering lectin specificity and the need to broadly characterize the interactomes of soluble lectins in native microbial communities.

## Acknowledgements

We would like to thank Luke Besse for project management and the Broad Institute’s Microbial ‘Omics Core and Genomics Platform for sample processing and sequencing data generation. We also thank Damian Plichta and Gleb Pishchany at the Broad Institute for helpful discussions. We are grateful to all study participants who provided samples and to Helena Lau and Kevin Shannon for clinical sample coordination for the PRISM cohort.

## Author contributions

Conceptualization: CRI, DAW, LLK

Methodology: RLM, CRI, RLW, DS, TG, SR, LTTN, DAW, MH, MW, AD, RK, SG

Investigation: RLM, CRI, RLW, DS, TG, SR, LTTN, DAW, MH, MW, AD, RK, SG

Visualization: RLM, CRI, RLW, MW

Funding acquisition: RLM, CRI, FER, HV, RJX, LLK

Project administration: HV, EJA, RJX, LLK Supervision: HV, EJA, RJX, LLK

Writing – original draft: RLM, CRI, RLW, DS, MH, HV, LLK

Writing – review & editing:

## Data and materials availability

Illumina sequencing data will be deposited into a public repository (pending). All other data are in the main paper or supplementary materials.

## Supplementary Materials

### Materials and Methods

#### Protein expression and purification

Cloning of full-length N-terminal strepII-tagged hItln1 into the pcDNA4/myc-HisA vector backbone (Life Technologies) was described previously (*14*). Human MBL (*MBL2*, NM_000242) gBlock DNA Fragment (Integrated DNA Technologies) was Gibson ligated into a linearized pcDNA4 plasmid (primers REV (5’-GGTGAAGCTTAAGTTTAAACG-3’) and FWD (5’-TGAGGATCCACTAGTCCAGTG-3’)). StrepII tag (5’-TGGAGCCATCCGCAGTTTGAAAAG-3’) was inserted C-terminal of amino acid 20 into pcDNA4-hMBL2 using inverse PCR mutagenesis using FWD primer (5’-TGGAGCCATCCGCAGTTTGAAAAGGAAACTGTGACCTGTGAGG-3’) and REV primer (5’-TTCTGAGTAAGACGCTGCC-3’). Correct insertion was verified by DNA sequencing (Quintara Biosciences).

hItln1 and MBL were each expressed by transient transfection of suspension-adapted HEK 293T cells. Cells were transfected at 1.8 x 10^6^ cells/mL in growth medium (DMEM (Thermo Fisher, cat. no. 11995) supplemented with 10% heat-inactivated FBS, 50 U/mL penicillin-streptomycin, 4 mM L-glutamine, and 1X non-essential amino acids) using lipofectamine 2000 (Thermo Fisher) following the manufacturer’s protocol. Six hours after the transfection, culture medium was exchanged to FreeStyle F17 expression medium (Thermo Fisher) supplemented with 50 U/mL penicillin-streptomycin, 4 mM L-glutamine, 1x nonessential amino acids, 0.1% heat-inactivated FBS, and 0.1% pluronic F-68 (Thermo Fisher). Transiently transfected cells were cultured up to 4 days or until viability was below 60%. The conditioned expression medium was then harvested by centrifugation and sterile filtration.

For purification, CaCl_2_ was added to the harvested conditioned expression medium to a final concentration of 10 mM, avidin (7 mg/mL) was added at 12 μL per mL of expression media (IBA, cat. no. 2-0204-015, per the IBA protocol). Protein was captured onto 2 mL of Strep-Tactin Superflow High-Capacity resin (IBA Lifesciences, cat. no. 2-1208-002) equilibrated with HEPES-Ca buffer (20 mM HEPES pH 7.4, 150 mM NaCl, 10 mM CaCl_2_). The resin was then washed with HEPES-EDTA buffer (20 mM HEPES pH 7.4, 150 mM NaCl, 1 mM EDTA), and eluted with 5 mM *d*-desthiobiotin (Sigma) in HEPES-EDTA buffer. StrepII-tagged hItln1 was concentrated with a 30,000-MWCO Amicon Ultra centrifugal filter. StrepII-tagged MBL was concentrated with a 10,000-MWCO Vivaspin 6 centrifugal filter (GE). All proteins were buffer exchanged to HEPES-EDTA buffer for storage. Protein concentrations were determined by absorbance at 280 nm. Extinction coefficients and molecular weights were calculated for the mature trimeric product of each protein (without the signal peptide) using the ProtParam tool (web.expasy.org/protparam). StrepII-tagged hItln1 had a calculated ε = 239,775 cm^-1^ M^-1^ and an estimated molecular mass of 102,024 Da. StrepII-tagged MBL had a calculated ε = 71,595 cm^-1^ M^-1^ and an estimated molecular mass of 75,570 Da.

#### Culturing of microbe isolates

Bacterial isolates along with their respective medias used in this study are detailed in Table S1 and Table S5. All isolates were struck out from frozen stock onto solid medium in an anaerobic chamber and incubated anaerobically at 37°C for 24-72 hours (isolate-dependent). Liquid Mega Medium (isolates in Table S1) was prepared and reduced in a Coy Laboratory anaerobic chamber (5% H_2,_ 20% CO_2_, 75% N_2_) as previously described (*42*). Liquid cultures were established for each isolate through the inoculation of a single colony into 5-10 mL of media. Upon the observation of turbid growth (16-72 hours), liquid cultures were combined with a 1x PBS + 40% glycerol + 0.001% L-cysteine solution at a 1:1 ratio, aliquoted into a series of cryovials, and stored at -80°C prior to flow cytometry. A portion of each culture was subjected to 16S rDNA V1-V9 PCR amplification, spin column purification, and Sanger sequencing, with taxonomic verification via standard nucleotide BLAST via blastn suite software (NIH, National Library of Medicine, https://blast.ncbi.nlm.nih.gov/Blast.cgi).

#### Flow cytometry of microbe isolates

All buffers were sterile filtered before use and all centrifugation steps were performed at 5000 RCF at 4°C. Microbe isolates were screened for viability by propidium iodide staining by flow cytometry prior to lectin binding. We observed differences in binding to viable and unviable cells; therefore, only viable isolates were used for validation. Microbe isolates washed 2X with 2mL PBS-BSA-T (PBS pH 7.4, Gibco, 0.1% (w/v) BSA (US Biological; A1311), 0.05% (v/v) Tween-20 (Sigma), pelleted, and resuspended in PBS-BSA-T. Staining was performed at OD_600_ of 0.2 for all samples. Staining solutions were HEPES-Ca-BSA-T with SYTO BC (1:1000, Thermo Fisher Scientific) and StrepMAB classic DY649-conjugate (1:150, IBA Lifesciences) for the unstained samples. All lectin-stained Ca^2+^ samples were stained as follows: HEPES-Ca-BSA-T, SytoBC (1:1000; Fig. 5, Fig. S5), StrepMAB classic Oyster 645 (1:250, Fig. 1, Table S1) or StrepMAB classic DY649-conjugate (1:250; Fig. 5, Fig. S5), 20 μg/mL recombinant StrepII-tagged lectin. For the lectin-stained EDTA control samples, the staining conditions were as follows: HEPES-EDTA-BSA-T (20 mM HEPES pH 7.4, 150 mM NaCl, 1 mM EDTA, 0.1% BSA, 0.05% Tween-20), SytoBC (1:1000), StrepMAB classic DY649-conjugate (1:250), and 20 μg/mL recombinant StrepII-tagged lectin. Staining was performed at 4 °C for one hour before being diluted 4X for analysis on a BD Accuri C6 flow cytometer (Fig. 1, Table S1) or ThermoFisher Scientific Attune flow cytometer (Fig. 5, Fig. S6).

#### hItln1 binding to synthetic mixed community

Strains of bacteria were prepared as described above. Cell density was quantified on a BD Accuri C6 flow cytometer and equal numbers of cells from each species were combined into a single sample. Cells were stained and analyzed as described above.

#### Direct labeling of proteins with fluorophores

Recombinant StrepII-tagged hItln1 and MBL were directly conjugated with fluorophores using NHS-ester conjugation chemistry. HItln1 was labeled with Alexa Fluor 647 NHS Ester (Succinimidyl Ester) (ThermoFisher Scientific) and MBL was labeled with Alexa Fluor 555 NHS Ester (Succinimidyl Ester) (ThermoFisher Scientific) following manufacturers protocol as follows: proteins were labeled in storage buffer (HEPES-EDTA, pH 7.4 as described above) by the addition of dye dissolved in DMSO at a ratio of 1:4 (w:w, dye:protein), and incubation for 1 hour at room temperature with mixing. Unreacted dye was removed by desalting using a Zeba 2 mL 7MWCO Spin Desalting Column (ThermoFisher Scientific) following manufacturer protocol. Molar ratio of fluorophore:protein was determined from UV-Vis absorbance spectrum using the extinction coefficients: ε Αlexa Fluor 555 = 155,000 cm^-1^ M^-1^, ε Alexa Fluor 647 = 270,000 cm^-1^ M^-1^.

### Glycan array analysis

Microbial glycan arrays were provided by Ryan McBride. For each lectin a microbial glycan array was thawed at room temperature for 30 minutes. The slide was placed in a slide holder tube and soaked in wash buffer (20 mM HEPES pH 7.5, 150 mM NaCl) for 10 minutes. The slide was placed onto a raised slide holder platform in a humidification chamber and a solution of StrepII-tagged lectin at a concentration of 10 μg/mL in binding buffer (20 mM HEPES pH 7.5, 150 mM NaCl, 10 mM CaCl_2_, 0.1% BSA, 0.1% Tween-20). The slide was incubated with gentle rocking at room temperature for 1 hour. The slide was then washed with binding buffer, wash buffer and ddH_2_O (4 times each). The slide was placed back into the humidification chamber and incubated with Dy549-conjugated anti-Strep Mab at 4 μg/mL in binding buffer at room temperature for 1 hour. The slide was washed again with binding buffer, wash buffer and ddH_2_O (4 times each). The slide was dried with a slide spinner and scanned using a Genepix4000 (Molecular Devices) using the 532 nm laser. Raw data was averaged across technical replicates and plotted against sample ID numbers.

### Approval for human sample research

Human stool samples were obtained from the Prospective Registry in IBD Study at MGH (PRISM) cohort, which was reviewed and approved by the Partners Human Research Committee (ref. 2004-P-001067). Stool samples from healthy subjects were obtained under protocol approved by the institutional review board at MIT (IRB protocol ID no. 1510271631).

Participants under both protocols provided informed consent and all experiments adhered to the regulations of the review boards.

### Preparation of human stool samples

Healthy human stool samples from IRB protocol ID no. 1510271631 were provided as a homogenized suspension by the Alm Lab (MIT). The preparation was as follows. Healthy subjects stool samples were brought into the anaerobic chamber within 2 hours of donation. Samples were homogenized using 1X PBS (pH 7.4), 0.1% L-cysteine at a ratio of 1 g stool:2.5 mL PBS. Glycerol (25% in 1X PBS, 0.1% L-cysteine) was added to the homogenate to a concentration of 12.5%, giving a final solution of 1g stool in 5mL 1X PBS, 0.1% L-cysteine, and 12.5% glycerol. The homogenate was removed from the anaerobic chamber and stored at -80°C prior to FACS analysis.

Fresh stool samples from adult donors with inflammatory bowel disease from the PRISM cohort were collected in anaerobic transport medium (Anaerobic Systems, AS-690), mailed to the Xavier Lab (Broad Institute) at ambient temperature, and processed anaerobically within 24-72 hours of collection. In the anaerobic chamber, samples were decanted into sterile gentleMACS C tubes (Miltenyi Biotec, 130-093-237) and topped up to a total volume of 10 mL using 1X PBS + 40% glycerol + 0.001% L-cysteine solution, respectively. Stool was homogenized via gentleMACS Dissociator (Miltenyi Biotec, 130-093-235) for 3 cycles of 61 s on the “intestine” setting, aliquoted into a series of 2.0 mL cryovials (Corning, 430659), and stored at -80°C prior to FACS analysis.

### Flow cytometry and FACS of donor stool samples

All buffers were sterile filtered before use and all centrifugation steps were performed at 5000 RCF at 4°C. Stool homogenate was thawed on ice and 200ul of homogenate was washed 2X with 2mL PBS-BSA-T (PBS pH 7.4, Gibco, 0.1% (w/v) BSA (US Biological; A1311), 0.05% (v/v) Tween-20 (Sigma), pelleted, and resuspended in PBS-BSA-T. The material was then passed through a 35 µm cell strainer cap (Falcon), pelleted, and resuspended in 2 mL HEPES-Ca-BSA-T (20 mM HEPES pH 7.4, 150 mM NaCl, 10 mM CaCl_2_, 0.1% BSA, 0.05% tween-20). A 5 μL sample of the bacterial suspension was pelleted and frozen as the input sample for metagenomic sequencing. Staining was performed at OD_600_ of 0.2 for all samples. Staining solutions were HEPES-Ca-BSA-T with SYTO BC (1:1000, Thermo Fisher Scientific) and StrepMAB classic DY649-conjugate (1:250, IBA Lifesciences) for the unstained samples. All lectin-stained Ca^2+^ samples were stained as follows: HEPES-Ca-BSA-T, SytoBC (1:1000), StrepMAB classic DY649-conjugate (1:250), 20 μg/mL recombinant StrepII-tagged lectin. For the lectin-stained EDTA control samples, the staining conditions were as follows: HEPES-EDTA-BSA-T (20 mM HEPES pH 7.4, 150 mM NaCl, 1 mM EDTA, 0.1% BSA, 0.05% Tween-20), SytoBC (1:1000), StrepMAB classic DY649-conjugate (1:250), and 20 μg/mL recombinant StrepII-tagged. Staining was performed at 4°C for four hours before being diluted 10X for analysis on a BD LSRII HTS flow cytometer with BD FACSDiva software, or 6X for FACS on a FACS Aria contained in a biosafety cabinet, with BD FACSDiva software. Stained samples were screened for viability by propidium iodide staining prior to lectin binding and sorting. Under our staining conditions lectin binding did not alter cell viability (Fig. S7). Cells were gated by forward scatter, side scatter, and SYTO BC staining, and sorted into lectin-negative or lectin-positive fractions. For metagenomics, 1.5x10^6^ cells were collected for the lectin-negative and - positive samples. Sorted cells were centrifuged at 7,000 RCF at 4°C for 5 minutes, supernatant was removed, and the cell pellet was stored at -20°C prior to nucleic acid extraction.

Flow cytometry on stool samples was performed at the The Swanson Biotechnology Center Flow Cytometry Facility housed in the Koch Institute (Massachusetts Institute of Technology, Cambridge, MA).

### Nucleic acid extraction

Cell pellets were lysed by addition of 50 μL HotShot Lysis buffer (25 mM NaOH, 0.2 mM EDTA, pH 12), and heating to 95°C for 10 minutes followed by addition of equal volume HotShot neutralization buffer (40 mM Tris-HCl, pH 5) and vortexed to combine. Lysed samples were centrifuged at 3000 RCF for 10 minutes to pellet debris and transferred to 96 well plates. The genomic DNA was then purified by AMPURE bead cleanup as follows: 90 μL beads was added to 100 μL DNA and incubated for 13 minutes at room temperature. Beads were separated on a magnet for 2 minutes and supernatant removed. Beads were washed 2X with 200 μL EtOH and air dried for 20 minutes while still on the magnet. The elution was performed by resuspending the beads thoroughly in 30 μL H_2_O and incubation for 7 minutes at room temperature. Beads were separated on a magnet and 27 μL eluted DNA was removed to a sterile plate.

### Metagenomic sequencing

#### Metagenomics Library Construction and Sequencing

Purified DNA was used as input (1 µL) into a miniaturized version of the Nextera-XT Library Preparation Kit (Illumina Inc.). All reactions were scaled to one-fourth their original volumes. Libraries were constructed according to the manufacturer’s instructions with several modifications to accommodate low DNA concentrations. The amplicon tagmentation mix (ATM) was diluted 1:10 in tagmentation DNA buffer (TD) to reduce the tagmentase:DNA ratio. Tagmentation time was also reduced from 5 minutes to 1 minute. Both modifications were implemented to boost the insert size thereby reducing read overlap and allowing for sampling of a larger proportion of the nucleotide sequence space. Lastly, the number of cycles in the library amplification PCR was increased from 12 to 20 to generate sufficient product for sequencing. Individual sample libraries were pooled at equimolar concentration and sequenced on a HiSeq 2500 at 200 cycles for 2x100 paired-end reads.

#### Metagenomic Profiling and Downstream Analysis

Raw sequencing reads were first processed through CutAdapt (version 1.7.1) (*43*) and Trimmomatic (*44*) (version 0.3.3) to remove Nextera adapters, low quality bases, and low quality reads. Trimmed reads were then filtered for human contamination (Hg19) with KneadData (version 0.5.1). These filtered reads were then run through Metaphlan2 (version 2.6.0) for taxonomic profiling (*31, 45*). Relative taxonomic abundances from Metaphlan2 were imported into the phyloseq package (version 1.30.0) for further refinement and visualization.

We categorized microbial species that were found in buffer controls as contaminants (most notably, the skin-resident microbe *Cutibactierum acnes*) and excluded these species from downstream analyses. Relative taxonomic abundances were used to calculate enrichment scores (Probability Ratios; P_R_) in R using the IgAScores package (version 0.1.2) (*32*). To calculate probability ratio enrichments in the case of a species present in the lectin-bound fraction but undetected in the input, we added a pseudo-count of 0.00001 for the abundance of that species in the input sample. Data were further processed using TidyR (version 1.1.4) and visualized by ggplot2 (version 3.3.5).

##### Supplementary Figures

**Figure S1.**
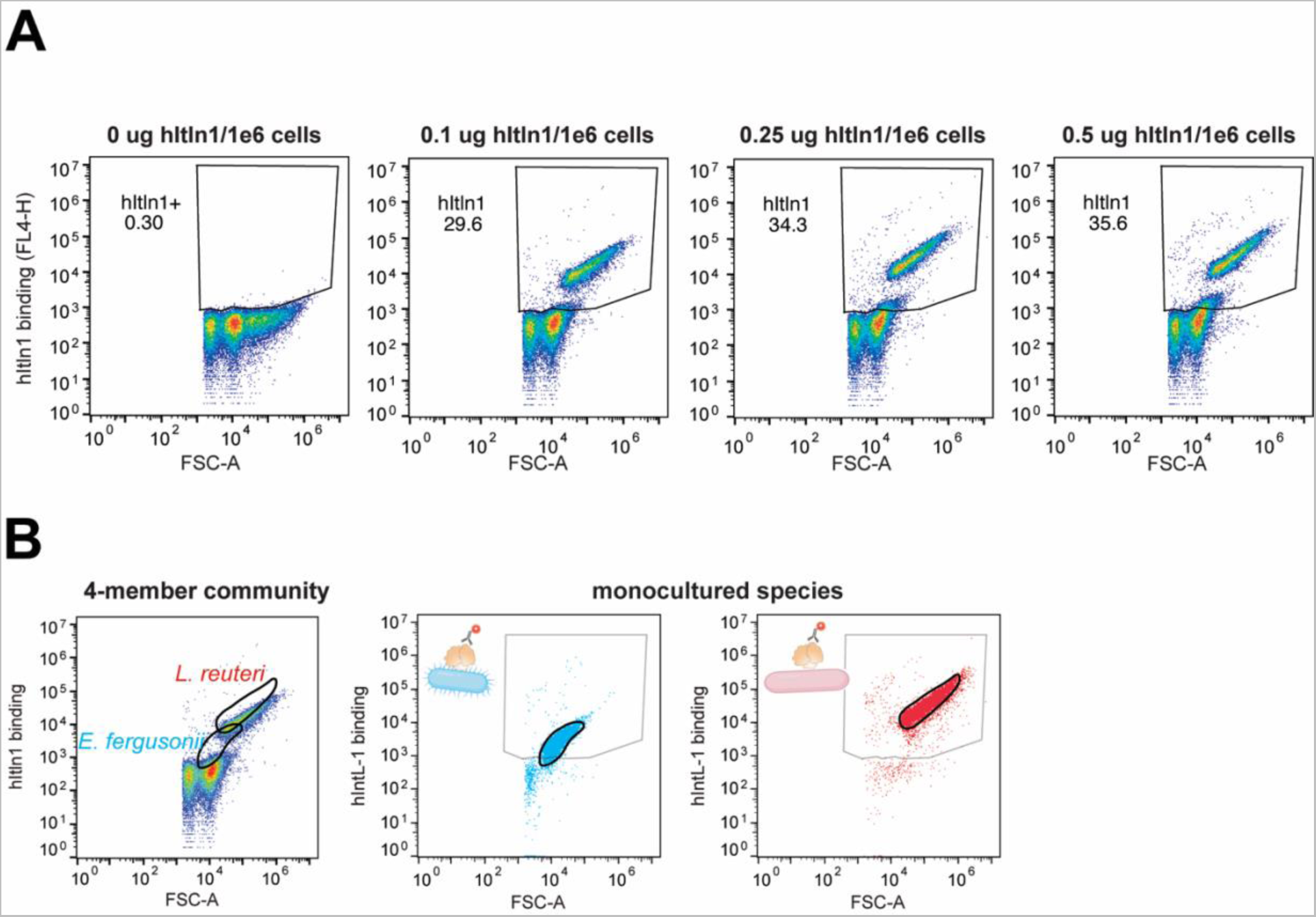
hItln1 binding to synthetic mixed microbial community. (A) Flow cytometry analysis of synthetic mixed microbial community plotted as forward scatter (FSC) against hItln1 binding (anti-Strep Oyster 645). Panels show from (-) lectin/Ca^2+^ and (+) hItln1/Ca^2+^ conditions with increasing concentrations of hItln1. (B) Flow cytometry analysis of synthetic mixed microbial community plotted as forward scatter (FSC) against hItln1 binding (anti-Strep Oyster 645) (data from Figure 1C) indicating the expected scattering patterns of *L. reuteri* and *E. fergusonii*. Panel shown is from 0.1 μg hItln1/1e6 cells condition. hItln1 binding to *L. reuteri* and *E. fergusonii* in monoculture binding assays are shown as reference (data from Figure 1B).

**Figure S2.**
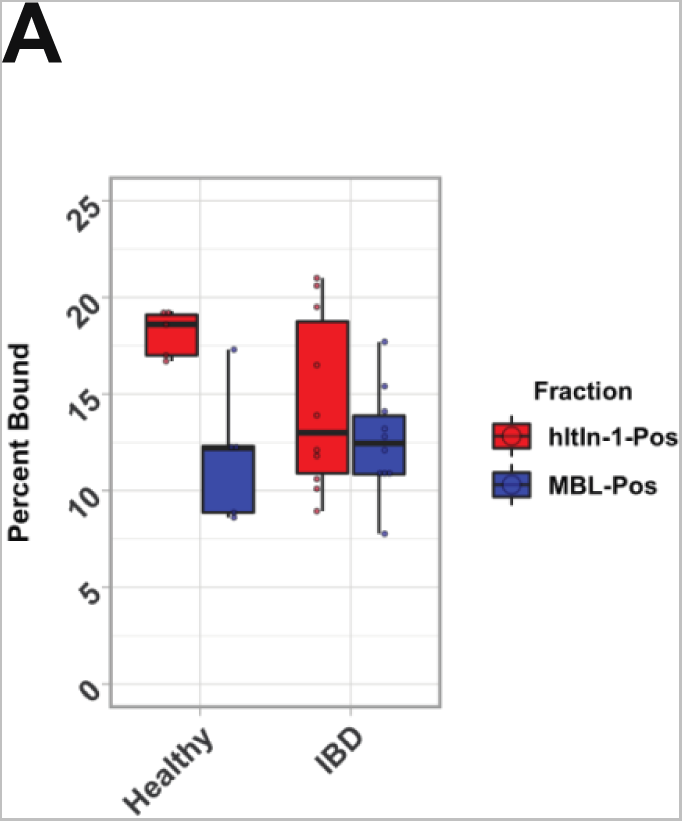
Binding of MBL and hItln1 to healthy or diseased human stool homogenate. (A) Quantification of binding of MBL and hItln1 to human stool homogenate collected from healthy or IBD-diagnosed donors. Percent bound is calculated by determining the percentage of cells that fall in the strep-pos/SytoBC-pos fraction (Q2). Data shown are from healthy donor samples bound by (+) strep-hItln1/Ca^2+^ (red; n=5; 18.1 +/- 0.5 % (mean, SEM)) and (+) strep-MBL/Ca^2+^ (blue; n = 5; 11.9 +/- 1.6 %) or IBD-diagnosed donor samples bound by (+) strep-hItln1/Ca^2+^ (red; n=9; 14.5 +/- 1.4 %) and (+) strep-MBL/Ca^2+^ (blue; n = 9; 12.6 +/- 0.9 %).

**Figure S3.**
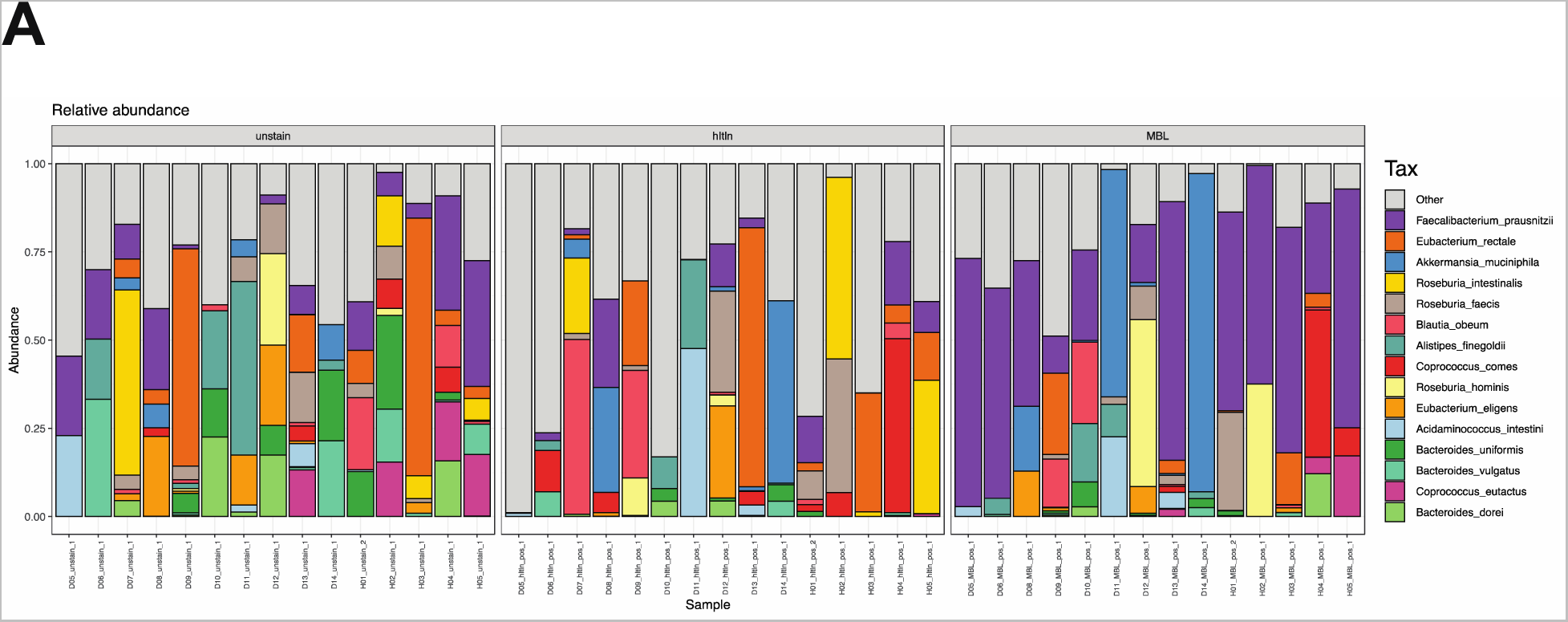
Fraction of bacterial species in input or lectin-bound fractions of human stool samples. Fractional abundance of bacterial species in the input (panel 1), MBL-bound (panel 2) or hItln1-bound (panel 3) fractions from human stool samples. Each column represents a different sample and species are represented by indicated colors.

**Figure S4.**
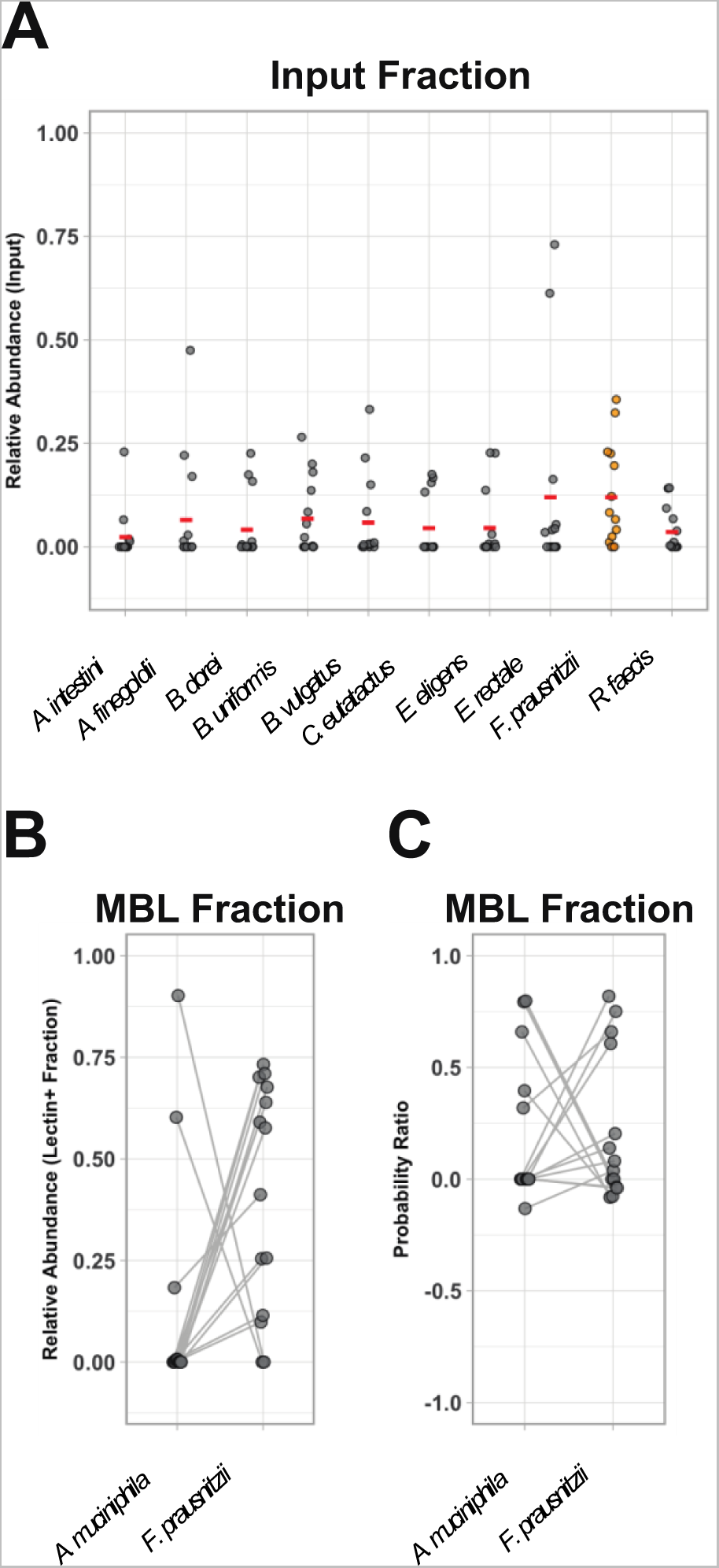
Abundance of bacterial species in input or MBL-bound fractions of human stool samples. (**A**) Dot plot of the relative abundance of top 10 most abundant species in the input fraction. (B-C) Relative abundances (B) or probability ratios (C) of *A. muciniphila* and *F. prausnitzii* in the lectin-bound fraction (MBL; all samples). Lines connect data points that come from the same sample.

**Figure S5.**
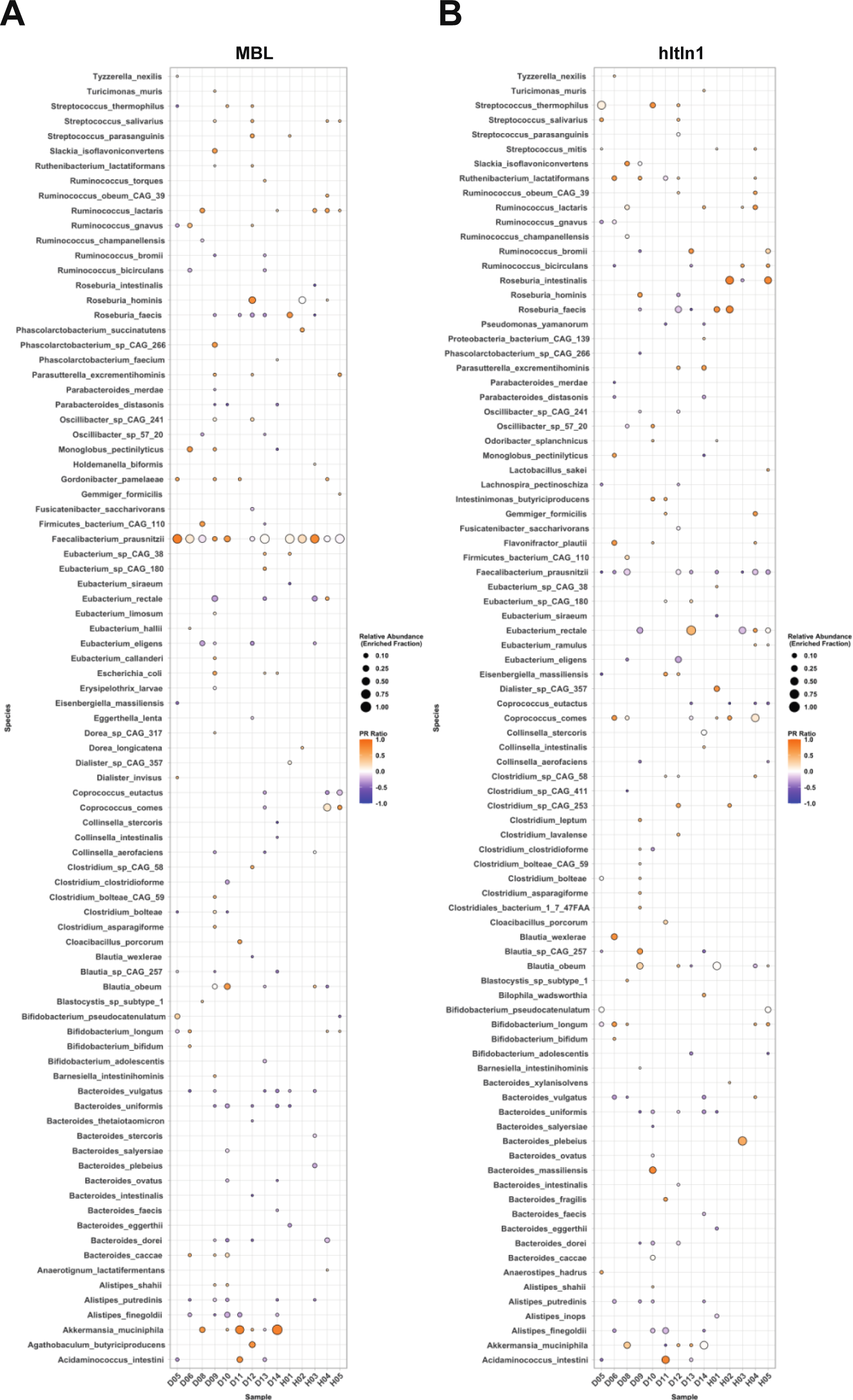
Probability ratios of MBL and hItln1 enrichments of bacterial species from human stool samples. “Bubble plot” depicting the enrichment of all enriched bacterial species in the lectin-bound fraction (defined as the Probability Ratio) donor stool samples bound by either MBL (A) or hItln1 (B).

**Figure S6.**
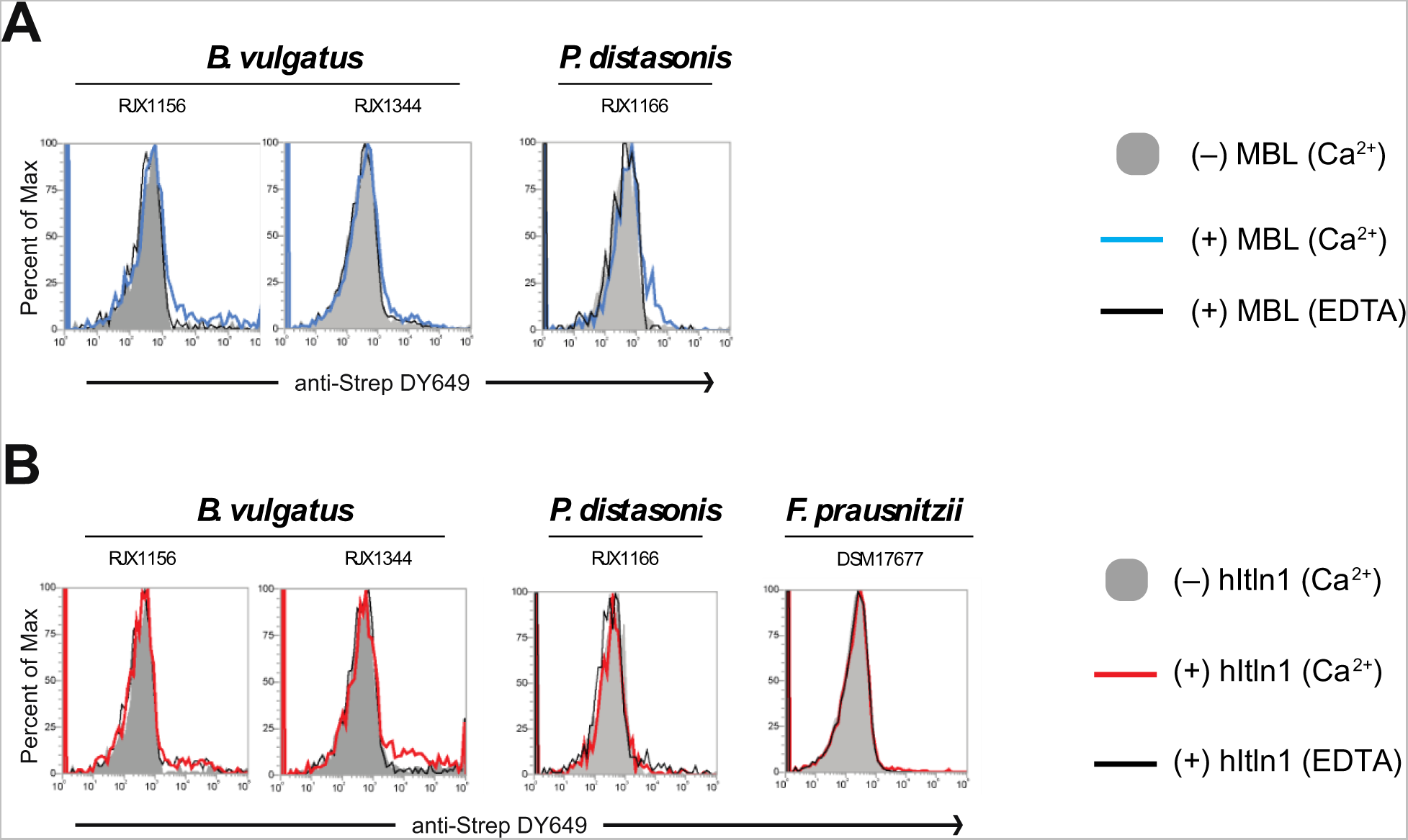
MBL- and hItln1-binding to negative control bacterial isolates. (**A-B**) Summarized flow cytometry analyses of MBL (A) or hItln1 (B) binding to bacterial isolates. Isolates are grouped by species. Individual histograms plot cell counts as a percent of the maximum signal against lectin binding (anti-Strep DY649). Each plot shows data from (-) lectin (Ca^2+^; solid grey), (+) MBL/Ca^2+^ (blue trace) or hItln1/Ca^2+^ (red trace), and (+) lectin/EDTA (black trace) binding conditions. Data are representative of two independent experiments.

**Figure S7.**
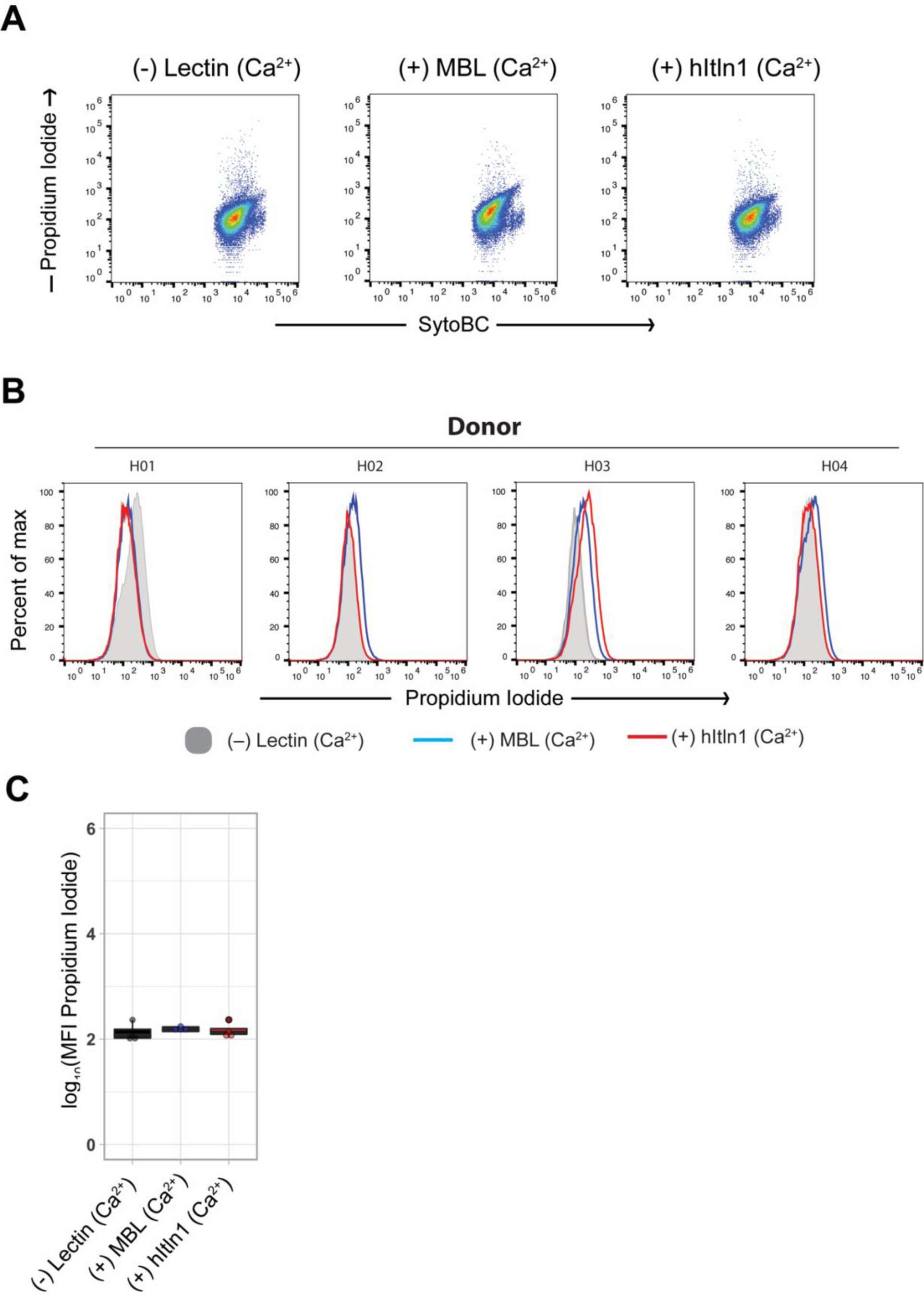
Effect of MBL and hItln1 binding on microbial cell viability. (**A)** Flow cytometry analysis of healthy human stool homogenate plotted as SytoBC against propidium iodide. Panels show no lectin control, (+) MBL (Ca^2+^) and (+) hItln1 (Ca^2+^) conditions. (B) Summarized flow cytometry analyses showing the effect of lectin binding on propidium iodide staining of healthy human stool homogenate samples. Individual histograms plot cell counts as a percent of the maximum signal against propidium iodide. Each plot shows data from (-) lectin (Ca^2+^; solid grey), (+) MBL/Ca^2+^ (blue trace) or hItln1/Ca^2+^ (red trace). (C) Quantification of data shown in A. Data shown are the mean fluorescence intensities of propidium iodide in the (-) lectin control (black; n=4; 135.3 +/- 30.1 (mean, SEM)), (+) MBL (Ca^2+^) (blue; n=4; 154.9 +/- 9.5) and (+) hItln1 (Ca^2+^) (red; n = 4; 145.6 +/- 27.8) samples.

##### Supplemental Tables

**Table S1.** Metadata and hItln1 binding summary of microbial isolates shown in Fig. 1.

**Table S2.** Raw glycan array data shown in Fig. 2D.

**Table S3.** Abundance and enrichment of microbial species in Lectin-Seq fractions.

**Table S4.** Stool sample metadata.

**Table S5.** Metadata of microbial isolates shown in Fig. 5 and Fig. S6

